# Trapped ion mobility spectrometry-guided molecular discrimination between plasmalogens and other ether lipids in lipidomics experiments

**DOI:** 10.1101/2024.10.23.619801

**Authors:** Jakob Koch, Lukas Neumann, Katharina Lackner, Monica L. Fernández-Quintero, Katrin Watschinger, Markus A. Keller

## Abstract

Trapped Ion Mobility Spectrometry (TIMS) has demonstrated promissing potential as a powerful discriminating method when coupled with mass spectrometry, enhancing the precision of feature annotation. Such a technique is particularly valuable for lipids, where a large number of isobaric but structurally distinct molecular species often coexist within the same sample matrix. In this study we explored the potential of ion mobility for ether lipid isomer differentiation. Mammalian ether phospholipids are characterized by a fatty alcohol residue at the *sn*-1 position of their glycerol backbone. Ether lipids can make up to 20% of the total phospholipid mass. They are present in a broad range of tissues where they are for example crucial for nervous system function, membrane homeostasis and inter-as well as intracellular signaling.

Molecular ether lipid species are difficult to distinguish analytically, as they occur as 1-*O*-alkyl and 1-*O*-alkenyl subclasses, with the latter being also known as plasmalogens. Isomeric ether lipid pairs can be separated with reversed-phase chromatography. However, their precise identification remains challenging due to the lack of clear internal reference points, inherent to the nature of lipid profiles and the lack of sufficient commercially available standard substances.

Here, we demonstrate - with focus on phosphatidylethanolamines - that ion mobility measurements allow to discriminate between the ether lipid subclasses through distinct differences in their gas phase geometries. This approach offers significant advantages as it does not depend on the variability of chromatography. However, the current resolution in the ion mobility dimension is not sufficient to baseline separate 1-*O*-alkyl and 1-*O*-alkenyl isobars, and the observed differences are not yet accurately represented in existing collision cross section databases.

Despite these challenges, the predictable properties of the ion mobility behavior of ether lipid species can significantly support their accurate annotation and hold promise for future advancements in lipid research.

## Introduction

Lipids are a diverse class of biomolecules and play crucial roles in a diverse range of cellular functions, including energy storage, membrane formation, regulation of membrane fluidity, as well as inter- and intra-cellular signalling^1–3^. Accurately characterizing lipid species is fundamental to comprehensively understand their biological roles and potential involvement in diseases. However, the respective method of choice - mass spectrometry (MS) - often faces challenges in differentiating isomers and resolving isobaric species with identical, as well as almost similar masses ^4^.

This is especially relevant for ether lipids, which can be subdivided into 1-*O*-alkyl and 1-*O*-alkenyl lipids, also known as plasmalogens (Figure 1). Ether lipids can represent up to 20% of all phospholipids (PL) in cells and tissues^5^ and play a critical role in various biological processes, for example in form of the 1-*O*-alkyl lipid platelet activating factor (PAF)^6^ or the 1-*O*-alkyl linked GPI anchors^7^. The functional role of the Δ1 vinyl ether double bond (DB) characteristic for plasmalogens is still largely unexplored^8^, also because of a lack of reliable analytical tools. Differentiating 1-*O*-alkenyl from their 1-*O*-alkyl counterparts remains a significant analytical challenge due to their identical sum formulas rendering them non-distinguishable in the mass dimension^9^.

**Figure 1:**
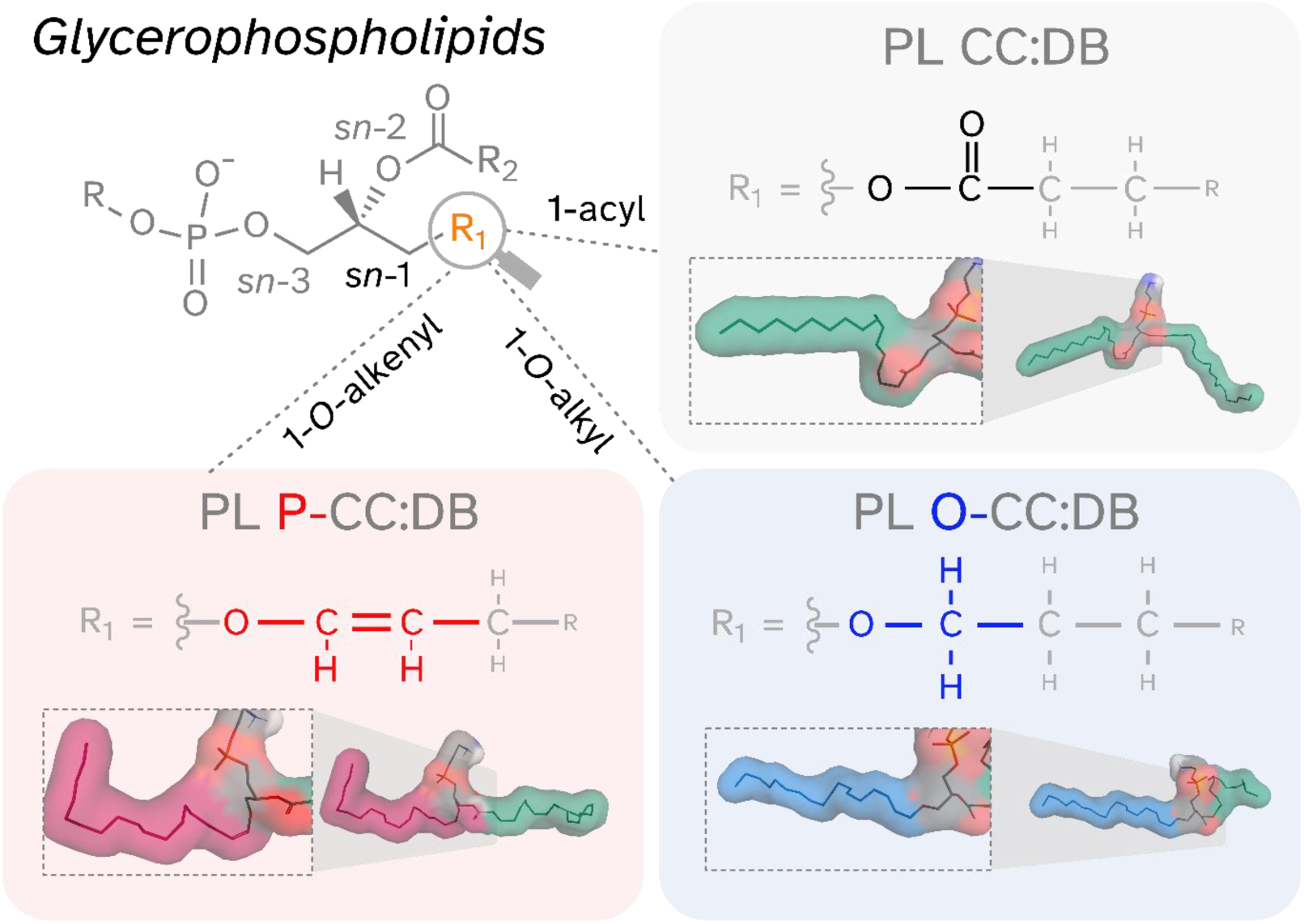
Core structures of fatty radyl linkage types of glycerophospholipids at the sn-1 position. Phospholipids (PL) consist of a glycerol backbone, a variable headgroup (R), the sn-2 fatty acyl chain (R_2_) and the sn-1 linkage site (R_1_, orange). 1-acyl (grey background, green sphere in structural representation), 1-O-alkyl (blue background and sphere), and 1-O-alkenyl (red backgroundwith pink sphere) linkages are shown along with one of the gas phase structures of the respective lipid molecule (red sphere color for O atoms). CC: total carbons in radyl side chains, DB: total number of double bonds in radyl side chains.

In this regard, ion mobility spectrometry (IMS) emerges as a powerful complementary technique, separating ions based on their size, shape, and charge in the gas phase. Indeed, the ether linkage type at the *sn*-1 position has been predicted to change the molecular surface geometry of the gaseous lipid molecule^10^. Such geometry differences result in different transition times in the presence of an inert gas stream throughout an IMS cell. Drift times can be converted to easier comparable collision cross section (CCS) values, which serve as an unique fingerprint for a specific ion under comparable conditions^11^. When applicable for the measurement in complex lipid mixtures, the integration of IMS with MS has thus - at least theoretically - the capacity to provide an additional layer of information, enhancing lipid characterization capabilities^12^.

Recent development of high-performance mass spectrometers suitable for omics studies with IMS capabilities has led to a renewed interest in 4D lipidomics^13,14^. Studies have explored the general applicability of IMS for lipidomics, culminating in the creation of valuable resources such as the lipidome-atlas and the CCS-compendium^15–18^. These databases provide a wealth of CCS data for different lipid classes, also allowing to predict the behavior of unknown lipid species. Additionally, detailed studies focusing on specific lipid molecules have further demonstrated the power of IMS for resolving complex lipid mixtures^19^. Hence, it has been shown, that lipid CCS are critical parameters for structural characterization and differentiation of lipids in ion mobility-mass spectrometry (IM-MS) based lipidomics workflows. However, despite their biological abundance, an explicit implementation of ether lipids and plasmalogens has so far been largely missed out. However, the need to accurately characterize plasmalogens and other ether lipids is underscored by their involvement in human diseases^20^, such as inherited peroxisomal disorders^21^, plasmalogen deficiency in the context of neurodegenerative diseases^22–25^ and cardiovascular diseases^5^.

The discovery of the genetic identity of plasmanylethanolamine desaturase 1 (PEDS1)^26–28^, the only known enzyme able to introduce the characteristic vinyl ether DB to form plasmalogens^29^, now opens up the possibility to explore the specific function of this DB in detail, independent of general ether lipid abundance effects^8^. Recently, we systematically analyzed the chromatographic reversed-phase separation behavior, which could be a potent approach to differentiate between isomeric ether lipid sub-classes. However, in mammalian tissue samples, the appearance of isobaric 1-*O*-alkyl and 1-*O*-alkenyl species is typically mutually exclusive, resulting in a lack of dataset-intrinsic reference points. Additionally, there is still a considerable lack of suitable commercially available isotope-labeled standard substances. This results in an urgent need for chromatography-independent strategies to achieve a reliable annotation of ether lipids, especially in untargeted lipidomics experiments.

In this study we make use of tissue material obtained from *Peds1*-deficient and wild type mice^26^, providing samples that either contain, or completely lack 1-*O*-alkenyl lipids^29^. This represents an ideal and physiologically-relevant model system to explore the power of IM-MS to differentiate between 1-*O*-alkyl and 1-*O*-alkenyl lipids. By combining the mass-resolving power of MS with the structural insights provided by IMS, and the separation capabilities of liquid chromatography (LC), we were able to accomplish an important step towards a fast and reliable characterization of these critically understudied lipid species in authentic biological sample matrixes.

## Results

A prerequisite for accurate discrimination between 1-*O*-alkyl and 1-*O*-alkenyl PL using IMS is that respective isomeric lipid pairs occupy distinguishable geometries in the gas phase. Thus, in a first step, we conducted force field geometry optimizations for representative pairs of 1-*O*-alkyl and 1-*O*-alkenyl lipids. Examples for such minimized structural solutions are shown in Figure 1. By comparing these predictions (Supplemental Table 1), as well as derived inverse mobility parameters (Supplemental Table 2), we could show that there are noticeable differences between the examined isomeric species (for more details see Supplemental Text 1). This demonstrated that ion mobility indeed has the theoretical potential to distinguish ether lipid pairs.

Next, we investigated the utility of IMS for identification and characterization of 1-*O*-alkyl and 1-*O*-alkenyl PL using mouse tissues as sample material. Particular focus was on phosphatidylethanolamine (PE) species, as distinguishing between radyl bonds is especially relevant within this group^29,30^. Mice were either wild type or *Peds1*-deficient (Δ*Peds1*), with the latter being characterized by a selective 1-*O*-alkenyl deficiency^29^. To cover a broad spectrum of different molecular lipid species, we chose heart, cerebellum, and cerebrum as model tissues, based on our previous experience with their ether lipid constitution^29,31^. Furthermore, male and female mice were considered. Mice were sacrificed, tissues harvested and homogenized, and lipids extracted. Then samples were subjected to LC-IMS-MS/MS measurement (see Methods for details). Special attention was on the optimization and calibration of the ion mobility range and trapped ion mobility spectrometry (TIMS) ramp settings (see Methods), using CCSbase^32^ as 1-acyl phospholipid reference for calibrating ether lipid CCS values in order to obtain high quality mobilograms. The annotation of molecular lipid species was performed up to the level of detail that the available data (retention time, exact mass, fragment spectra) allowed^33^. Figure 2A shows the PE O/P-38 series in detail and illustrates the observed ion mobility behavior of respective lipid species in a representative heart tissue sample of wild type (left panel) and *Peds1*-deficient (right panel) mice, respectively. As expected, PE ether lipids were found to be present as 1-*O*-alkenyl species in wild type and as 1-*O*-alkyl species in *Peds1*-KO mice. Characteristic DB number-related shifts in the m/z and retention time (RT) dimensions were detected (Figure 2B). Similarly, also in the ion mobility dimension (Figure 2A) we measured characteristic shifts for different lipid species, although with the limitation of a comparatively lower resolution.

**Figure 2:**
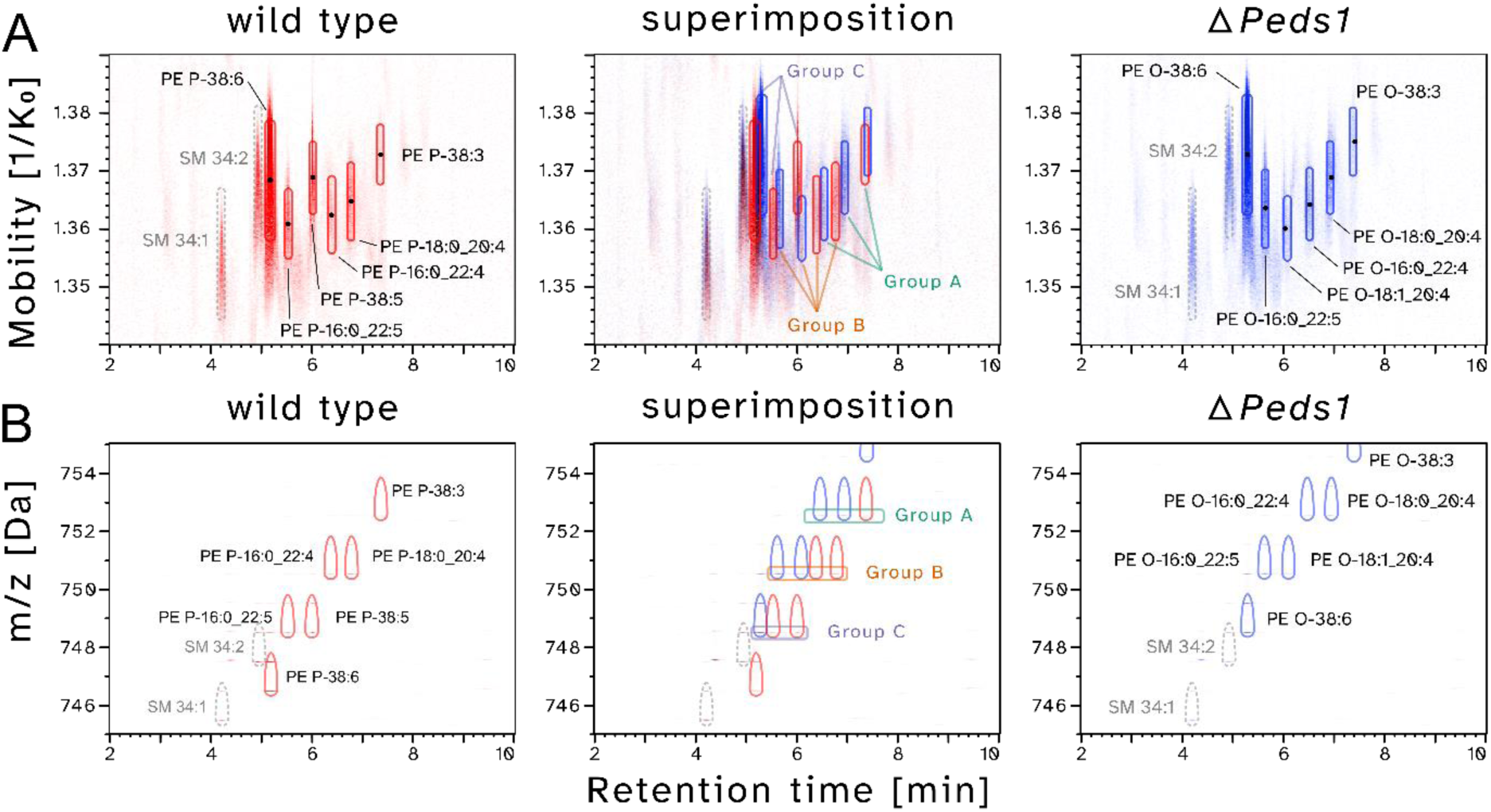
2D elution plots depicting the inverse ion mobility with m/z behavior of PE O/P-38:DB ether lipids. A) Retention times (in minutes) plotted against inverse ion mobility for wild type (red, left panel) and Peds1-deficient (blue, right panel, ΔPeds1) heart samples, as well as their superimposition (center panel). Black dots indicate the respective average measured inverse ion mobility values determined for reliably annotated PE features. Superimposition of similar intensities is indicated in black. Annotation of PE P-38:5 was partially ambiguous (see Supplementary Figure 1). B) Representation of retention times versus m/z ratio for the same samples shown in A). Please note that the boxes in the superimposition correspond to the [M-H]^−^ feature (m/z dimension), which was evaluated for IMS (panel A). Lipid abbreviations are in accordance with^33^, giving annotations on a lipid species and molecular lipid species level where applicable.

While the RT was strongly influenced by the cumulative number of DBs, the IMS readout additionally was found to be responsive to the exact molecular composition of lipid species. This resulted in a more complex IMS pattern, particularly at higher DB numbers (>5).

When analyzing the IMS behavior within groups of isomeric ether lipids, the exact quantitative differences of ion mobility values vary on a case-by-case basis (compare Groups A, B and C in Figure 2A). Although 1-*O*-alkyl species typically exhibit lower mobility values compared to their 1-*O*-alkenyl counterparts (Groups A and B), this is not necessarily true in every instance (compare Group C). When extending this analysis from the initial focus on the PE O/P-38 series to the entirety of reliably measurable PE species, it could be deduced that the previously described relationships are generalizable, as shown in a superposition of individual data points from six biological replicates (Figure 3). We observed the characteristic wing-shaped elution profile of PL on reversed-phase columns^34,35^ (Figure 3A). While the RT and m/z dimensions were highly reproducible, the individual ion mobility values (CCS [Å²], shown on the y-axis in Figure 3B and C) exhibited more variability.

**Figure 3:**
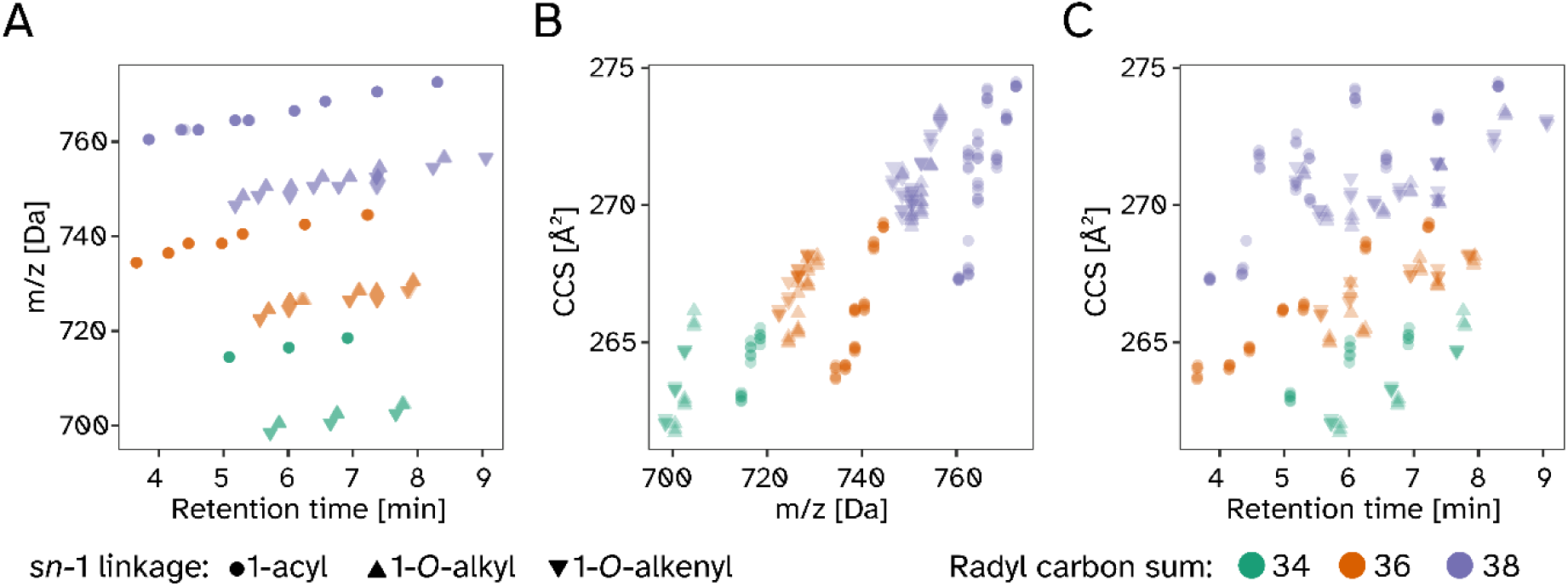
The extracted and averaged values for retention time, m/z, and collision cross section (CCS) of wild type and Peds1-deficient mouse heart homogenates in three replicates each. A) Retention time versus m/z plane, B) m/z versus CCS, C) retention time versus CCS. Shapes: 1-acyl (circles), 1-O-alkenyl (downwards pointing triangles), and 1-O-alkyl (upwards pointing triangles) colored according to the cumulative number of carbon atoms in the fatty radyl side chains: 34 (green), 36 (orange), 38 (purple).

This was particularly true for the m/z range above 720 Da, where a greater diversity of PE lipid species can be expected. There was also a positive correlation between m/z and CCS (Figure 3B). The relationships between the other dimensions are structured as well but followed a more complex pattern (Figure 3A and C).

While CCS was also influenced by the total number of double bonds, it was not just their presence but also their distribution and configuration within the respective molecule that determined the precise ion mobility behaviour. This explains the ordered and consistent CCS patters observed for lower molecular mass PE species (e.g., those with 34 cumulative carbon atoms in the radyl side chains (CC), Figure 3B) and the increased profile complexity seen in higher masses, which carry more double bonds (38 CC, Figure 3B). This effect was not limited to 1-*O*-alkyl / 1-*O*-alkenyl lipids but was also present in 1-acyl species. Furthermore, we also observed the same principal behavior also when analysing lipid extracts of cerebrum (Supplementary Figure 2) and cerebellum (Supplementary Figure 3) tissues of wild type and *Peds1*-deficient mice, however based on a different set of tissue-specific lipid side chain distribution patterns.

Figure 4A depicts characteristic examples for ion mobility traces of two isomeric 1-*O*-alkyl (blue) and two 1-*O*-alkenyl (red) lipid species in heart tissue. Ether lipids with longer *sn-*2 linked acyl side chains (22 CC) had lower CCS values compared to their shorter (20 CC) isomers and 1-*O*-alkyl had lower CCS values than respective isomeric 1-*O*-alkenyl species. As indicated by the vertical dashed lines (Figure 4A), each molecular species was characterized by a specific CCS value, which could, in principle, be used for their discrimination. At the same time, it was apparent that the CCS distributions of the individual isomeric species still significantly overlapped at the IMS resolution that could be achieved in this study. Thus, if species were to elute at similar RT, distinguishing them based on CCS alone would be particularly error prone.

**Figure 4:**
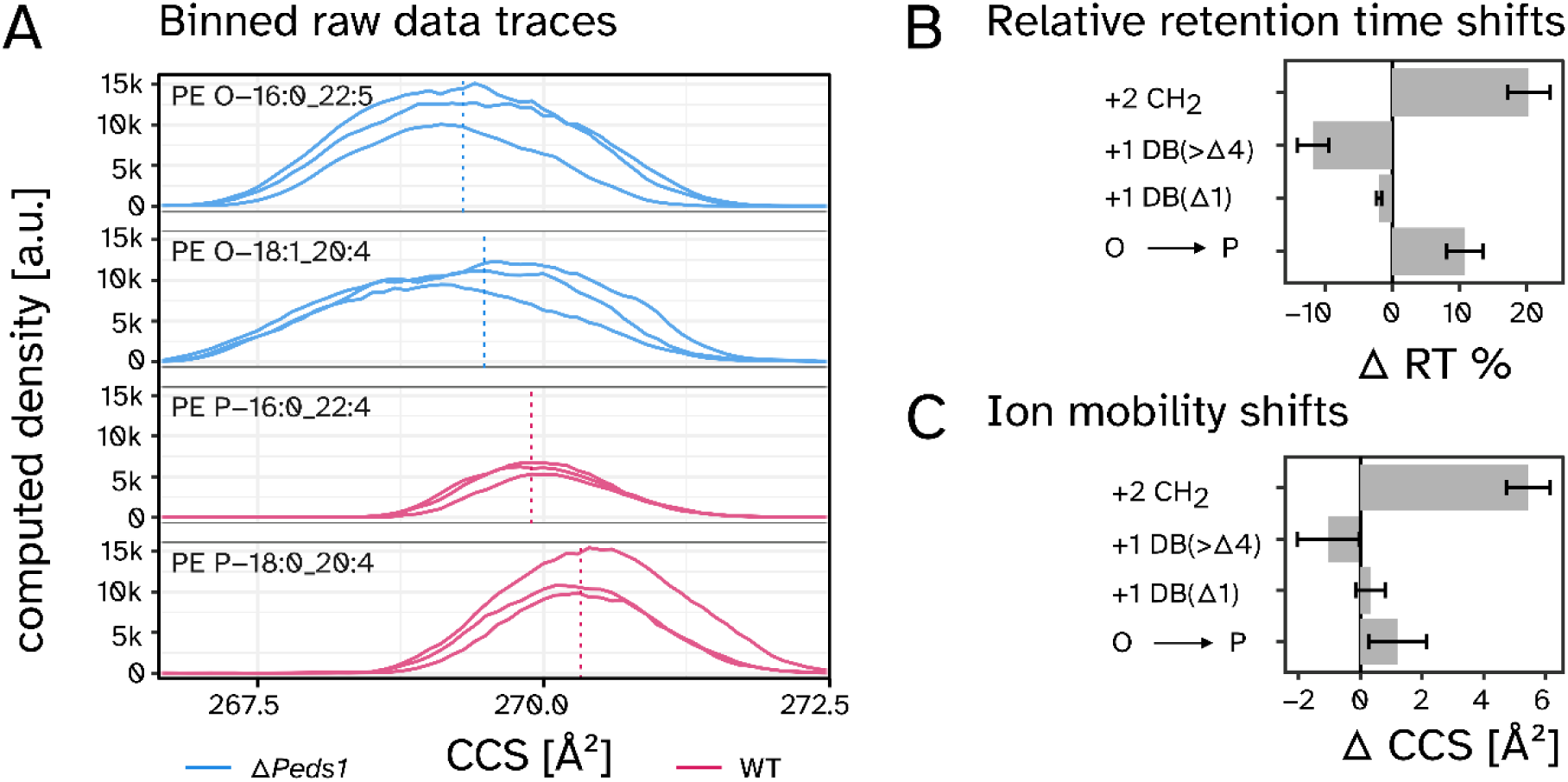
RAW data CCS traces and statistical assessment of influencing factors. A) Averaged CCS traces of representative ether lipid features in heart tissue samples (n = 3). The vertical dashed lines indicated the center of intensity per feature. B) Analysis of the impact of radyl chain elongation (+2 CH2), additional double bonds (+1 DB (>Δ4)), presence of a vinyl ether double bond (+1 DB (Δ1)), and the differential impact of 1-O-alkenyl and 1-O-alkyl lipids on retention times. Averaged relative differences for 10-32 relevant lipid species pairs each are shown. C) Analogue to B) but for the respective CCS values. Data shown as mean values ± SD. Analysis was based on lipid species from heart, cerebellum, and cerebrum of Peds1-deficient and wild type mice (n = 3 per group) with 10-32 comparisons on lipid species level.

In a next step, we systematically investigated the RT and CCS behavior of ether lipids, conducting between 10 and 32 individual comparisons of isomeric lipid pairs per parameter. We focused on the influence of an acyl chain extension by two CC, the impact of additional DB within the fatty radyl chain, and the specific contribution of plasmalogen double bonds. This analysis was previously performed for RT^29^, and showed highly similar results (Figure 4B). Crucial for distinguishing between 1-*O*-alkyl and 1-*O*-alkenyl species was the significant difference in RT depending on whether a double bond was located at the Δ1 position (i.e. in plasmalogens) (−2.0±0.4%) or further downstream in the radyl side chains (−11.7±2.3%). This resulted in an overall difference of 10.8±2.7% with the specific chromatographic system used in this study (Figure 4B).

In the CCS dimension, we observed changes comparable to those seen in RT (Figure 4C). CC was the most influential factor (+5.4±0.7 Å² for every two carbons), followed by DB (−1.0 ±1.0 Å² per DB). With a vinyl-ether DB shifting the CCS value by 0.3±0.5 Å² the overall net difference between 1-*O-*alkyl and 1-*O*-alkenyl was 1.2±0.9 Å² (Figure 4C). The relative effect size was therefore smaller than in the RT dimension. At the same time, the measured variance was higher, which was likely caused by the previously mentioned CCS differences between individual molecular species at higher numbers of DB.

As different databases of mostly computationally-derived CCS values were recently published, we next evaluated the performance of LipidCCS, a popular *in silico* CCS prediction tool^36^ (available at https://www.metabolomics-shanghai.org/LipidCCS/), by comparing its predictions with our experimentally measured CCS values (calibrated via 1-acyl PE obtained from CCSbase^32^) of phosphatidylethanolamine (PE) lipids encompassing 1-acyl, 1-*O*-alkyl, and 1-*O*-alkenyls. Linear regression analysis of measured vs. predicted CCS values revealed a perfect slope of 1.01 for 1-acyl lipids compared with 1.09 and 1.17 for 1-*O*-alkyl and 1-*O*-alkenyl lipids, respectively (Figure 5A). This indicates that, while 1-acyl are very well represented in the mathematical model based on LipidCCS database results (Supplementary Figure 5), there are still considerable room for improvement in the predictive power for 1-*O*-alkyl and especially 1-*O*-alkenyl lipids. A similar conclusion can be drawn from the Akaike Information Criterion (AIC), which measures how much information is lost by a model relative to other models. The high value for 1-acyl lipids indicates that here the LipidCCS model performs approximately two-fold better compared to lipids harboring one of the two ether lipid bondage variants at *sn*-1 (Figure 5B).

**Figure 5:**
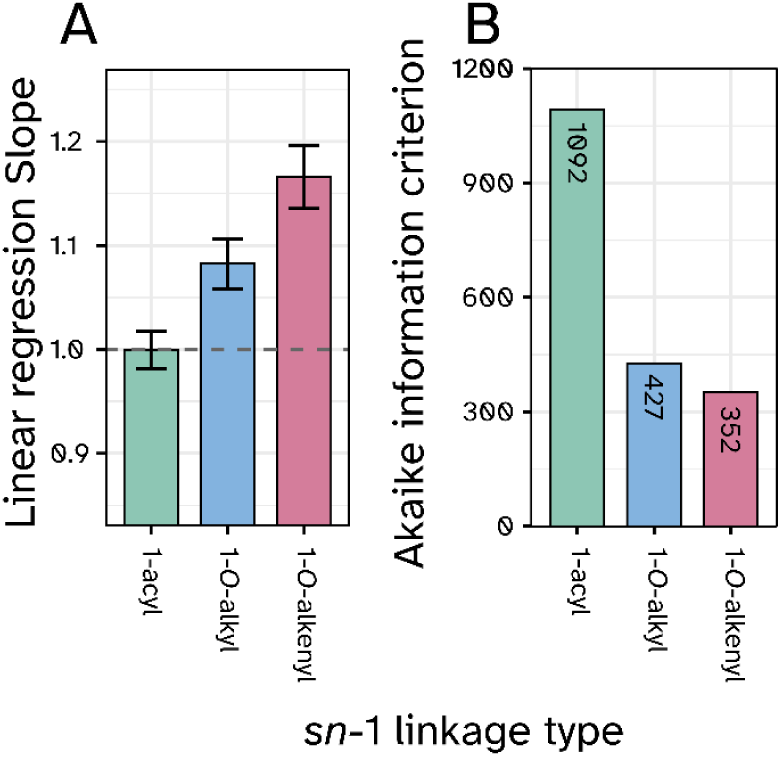
Comparison of measured and LipidCCS predicted values for phosphatidylethanolamine (PE) lipids stratified by subclass. A) Slopes of linear regression analysis between measured CCS values versus predicted CCS values for 1-acyl, 1-O-alkyl, and 1-O-alkenyl PE lipid subclasses as deposited in the lipidCCS database^36^. A slope of 1 indicates the best possible agreement (mean ± SD, n(1-acyl)=342, n(1-O-alkyl)=156, n(1-O-alkenyl)=126). B) Akaike information criterion (AIC) values for the comparison in A). Lower AIC values indicate worse model performance for the respective subclasses. All molecular lipid species in the database were averaged on lipid species level before being compared to their measured lipid species level counterparts.

Although the LipidCCS model turned out to be not entirely accurate for ether lipids, it still performed exceptionally well compared to predicting CCS values via proximity structure approximation (PSA) from minimized gas phase structures^37^. Despite a certain degree of linear relationship with the measured values, there was little quantitative agreement with the measurement results (further discussed in Supplementary Text 1).

To test to which extent the observed patterns can be explained mathematically, we constructed a linear model to predict RT and CCS values for individual lipid species. The model was based on characteristics we previously observed to be potentially influential. One example parameter is the cumulative DB number that generates highly predictable patterns in reversed-phase RT values (Figure 6A, upper panel). A comparable trend is also apparent in the CCS dimension, however with bimodal deviations at higher double bond numbers (Figure 6A, lower panel; see Supplementary Figure 6 for further subclasses). In this approach the cumulative numbers of double bonds and carbon atoms and the *sn*-1 binding types presented as the determining factors for observed RT and CCS behavior (Figure 6B). We tested linear models and also included bilinear interactions. All the parameters of the models under consideration are different from zero with statistical significance. Due to the categorical nature of the parameters in the end we consider a rather simple model with only 5 coefficients the best compromise between prediction quality and risk of overfitting (Result in Figure 6C).

**Figure 6:**
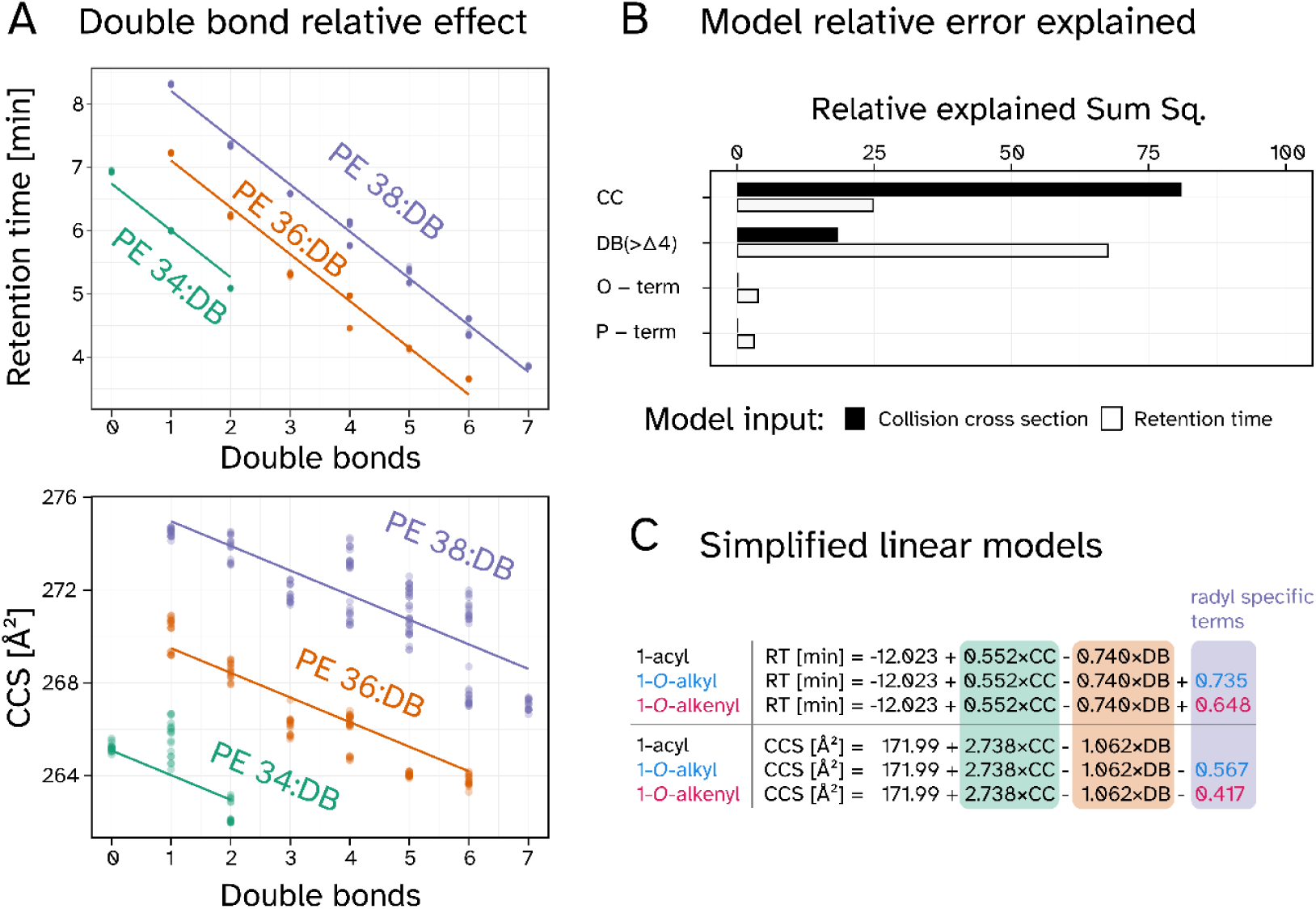
Linear combination models predicting 1-acyl, 1-O-alkyl, and 1-O-alkenyl retention times (RT) and CCS values. A) Link between double bond number and RT (upper panel) and CCS values (lower panel) stratified by carbon chain length. The 1-acyl subset of PE lipids was annotated according to cumulative side chain carbons (CC) in green (34), orange (36) and purple (38). Individual averaged values per sample and species were assessed from heart, cerebellum, and cerebrum tissues of Peds1-deficient and wild type mice (n = 3 per group). B) Relative sum of squares representing the errors of the linear combination models plotted for RT (red) and CCS (blue). Linkage type specific model errors are attributed as O-term (1-O-alkyl), and P-term (1-O-alkenyl). C: Linear equations derived from the models in respect to RT and CCS. Highlighted terms correspond to CC (green), total sum of double bonds in radyl-chains (DB, orange), and a variable ether lipid term (purple). Model contribution of ether lipids are indicated by text color blue (1-O-alkyl) and red (1-O-alkenyl).

Overall, the residual error showed that a significantly larger fraction of the information contained in the measurement could be explained in the RT dimension, compared to the CCS dimension (total sum of squares 868 vs. 6991, respectively). The same was also true for more complex models including more parameters and bilinear interactions. The carbon chain length (CC) was found to be most influential for CCS behavior, while RT was more strongly driven by the total number of DBs (Figure 6B). Both, CCS and RT are only marginally influenced by a contribution of an 1-*O*-alkyl or 1-*O*-alkenyl modifier, which represents an important distinguishing feature compared to the influence of other double bonds. From this overall behavior, a simple mathematical relationship could be derived, which can predict these parameters for any PE lipid species (Figure 6C).

To evaluate the power of ion mobility for ether lipid separation, we next determined the average ΔCCS values for 1-*O-*alkyl/1-*O*-alkenyl pairs with identical sum formulas that are indistinguishable by their mass to charge ratio alone. Out of the recorded ether lipid signals, 17 pairs qualified for such a direct comparison (see Supplementary Figure 7 for representative examples). Assuming Gaussian peak shapes for ion mobility - which is a necessary simplification for some features in mammalian tissue lipid extracts - we computed peak overlaps for 1:1 mixtures of these isomers at different ion mobility resolutions (Figure 7). This data was also used to calculate the respective theoretical resolution, as well as the resolving power respective to a CCS value of 268.4 Å² (average ether lipid in our dataset).

**Figure 7:**
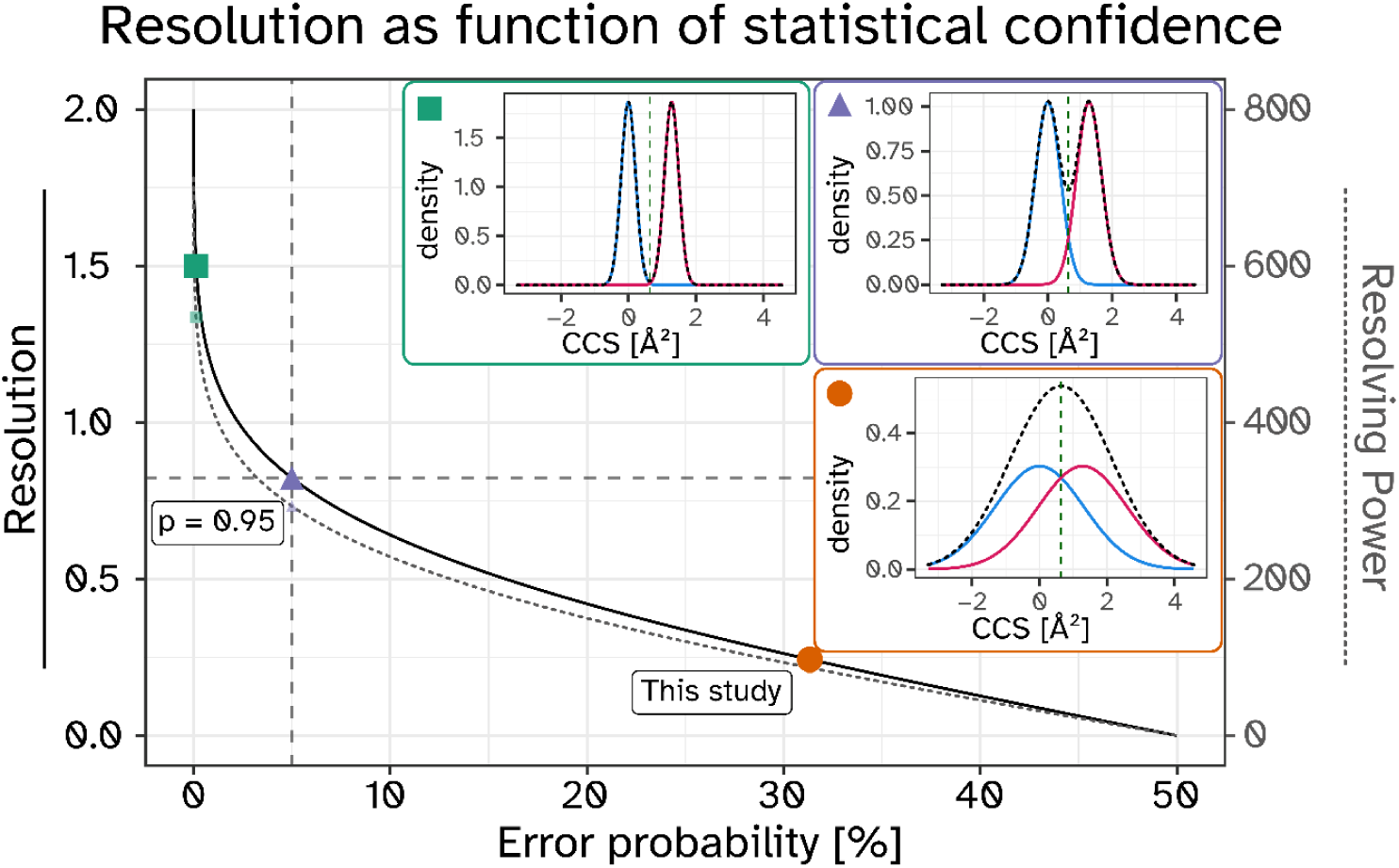
IMS resolution as a function of error probability when used as the sole criterion for isobaric ether lipid annotation. Orange circle and insert (fitted Gaussian models) indicate the average ether lipid resolution measured in this study (31% error probability). Similar distributions are shown for the scenarios of 5% residual annotation error (purple triangle, p = 0.95, R = 0.82) and baseline separation (green square, R = 1.5, p = 0.9987). For all 3 inserts the blue trace is positioned at µ = 0, and the second Gaussian trace (red) is placed at the average 1-O-alkyl/1-O-alkenyl difference observed in our raw data (1.28 Å²). The sum of both traces per subfigure is depicted as black dashed line. CCS-calibrated RAW data points were filtered for 1-O-alkyl and 1-O-alkenyl lipids, and an overall weighted standard deviation calculated with the r-function descriptio::weighted.sd().

With the analytical platform used in this study, we observed an average IMS resolving power of 87 (resolution=0.24), which translates into an error probability of 31% when annotating ether lipids purely on basis of their m/z and CCS properties (Figure 7, orange insert). A reduction of the error probability to 5% would be achieved when increasing the IMS resolving power to 293 (resolution = 0.82, purple insert). Full baseline separation and a corresponding error probability of 0.13% would be reached at a resolving power of 534 (resolution=1.5, green insert).

## Discussion

In this study, we employed a Trapped Ion Mobility Spectrometry (TIMS) capable instrument to investigate the behavior of the mammalian ether lipid variants of PL, with a specific focus on the potential of ion mobility for isomer differentiation. Ether lipids are present in almost all mammalian tissues, comprising up to approximately 20% of the phospholipid mass^31,38,39^. Furthermore, these lipids, particularly plasmalogens, play a crucial role in neurological health^40,41^, being abundant in the brain and contributing to neuronal communication^42,43^ and protection against oxidative stress^44,45^. Despite their importance, the functional differences between 1-*O*-alkyl and 1-*O*-alkenyl subspecies remain unclear^8^. This is to a large degree caused by the analytical challenges in accurately identifying them in mass spectrometric experiments ^46–48^.

### Instrument parameter and measurement principles

The focus on PE species in this study results from several factors. Firstly, this lipid class represented with the largest fraction of ether lipids in most used model systems and tissues ^29,38^. Secondly, the interplay between 1-O-alkyl and 1-O-alkenyl subspecies is most influential within PEs. This is due to the substrate specificity of PEDS1, which accepts 1-O-alkyl PEs. In contrast, Phosphatidylcholine ether lipids are less abundant, do not commonly carry a vinyl ether double bond, and are therefore not as severely impacted by a PEDS1-deficiency^29^. Furthermore, PE lipids exhibit a structural side chain diversity that is broad enough to enable the reliable annotation of a sufficient number of 1-O-alkyl and 1-O-alkenyl pairs for statistical comparisons. By selection of brain and heart tissues, we aimed to capture a representative profile of PE ether lipid species present across different mammalian tissues. As a result of this focus, we chose instrument parameters that were specifically optimized for the m/z and IMS range of PE ether lipids, which may not necessarily be the ideal settings for other lipid classes. In particular, the IMS range was optimized within a relatively narrow window to achieve the best possible resolution.

Accurate lipid annotations are critical for data interpretation^49–51^, especially given the vast amount of data generated per sample run^52^. We implemented this within our analysis and the structural description of lipid species reflects the information accessible within the available raw data. While all features exhibited consistent and robust RT, the IMS dimension showed broader variability due to the design and measurement principles of the trapped ion mobility instrument. This broader distribution required noise filtered averaging to allow for a reliable determination of mean ion mobility values^53^.

### Challenges and limitations of ether lipid species differentiation

While the m/z dimension offered superior peak sharpness and resolution for lipid identification, it lacked the ability to univocally distinguish isomers like 1-*O*-alkyl and 1-*O*-alkenyl PEs^29^. Although such a differentiation can be based on RT^29,54^, the exact chromatographic behavior is not easily transferable between different instrumental setups due to the heterogeneity of methods in use^55^. Even the same HPLC method run on different instruments can lead to RT changes, as is the case in this study, compared to earlier work of our group^29^. Additionally, it is challenging that equimolar mixtures of 1-*O*-alkyl and 1-*O*-alkenyl lipids are rare in biological samples. Typically, one species quantitatively dominates, which greatly increases the risk for misidentification during annotation.

In contrast, IMS offers a chromatography-independent approach for isomeric ether lipid differentiation in most cases, although the currently achievable resolution is not yet sufficient for baseline separation (Figure 7). However, when combined with chromatography in untargeted lipidomics experiments, IMS has the power to effectively aid the differentiation between 1-*O*-alkyl and 1-*O*-alkenyl lipids, thereby reducing the need for precise in-house RT databases (which are of course still highly recommended) and adding an additional layer of confidence in the annotation. Given the well-known lack of sufficient commercially available ether lipid standards, this technology could make an important contribution to improving the accurate representation of ether lipids in lipidomic data when utilized to its current full potential.

A possible limitation of this study was that Gaussian peak shapes were assumed for CCS traces (Figure 4), which was in some cases a necessary simplification for downstream analyses. Since our experiments were based on authentic lipid extracts rather than chemically pure substances, closely related non-ether lipid double bond isomers with similar or identical RT co-eluted in the IMS dimension. This contributed to broadening of CCS peaks. On larger scales, such as the differentiation between lipid classes^56,15^, total chain lengths^57,58^, and double bond numbers, these effects play a minor role. However, they did significantly contribute to shaping the raw mobilograms that might be useful for the finer structural elucidation of lipid species.

### Influence of fatty acyl substitution patterns on CCS and RT

It is a well-described phenomenon that the cumulative number of double bonds in a PE lipid strongly influences its RT, but to a much smaller extent their exact distribution between the two fatty acids^55,59^. Interestingly, a clearly more complex pattern was observed for IMS, particularly when lipids were carrying higher numbers of double bonds (Figure 6A). Deviations from a purely linear behavior were mostly driven by poly-unsaturated fatty acyl residues (FA) such as FA20:4 und FA22:6 (see Supplementary Figure 6). This finding aligns with earlier reports on the influence of fatty acyl structure on CCS^10,60,61^. With the present set of naturally occurring PE species, it is not entirely clearly assignable to what extent the differences were caused by the double bonds themselves or by the resulting different distribution of carbons atoms between the sn-1 and sn-2 positions. As previously shown for cardiolipins, raw data deconvolution could help to obtain the semi-quantitative contributions of individual molecular lipid species^62^. The exact influence of specific DB positions and distribution on the ion mobility behavior requires additional systematic investigation^63,53,64^ and should be examined in future studies independently of the here studied 1-O-alkyl and 1-O-alkenyl distinction.

### Importance of accurate annotation and database

Precisely validated lipid databases for exact mass, CCS, fragment spectra, and RT are pivotal for the field of lipid research. Respective data collections can be based on direct measurements^15,65^, as well as on computational strategies^32,36^. Comparison of the here determined CCS values with the LipidCCS database^36^ revealed an almost perfect agreement for 1-acyl PEs. In contrast, we observed discrepancies in the exact accordance for 1-O-alkyl and 1-O-alkenyl lipids (Figure 5), suggesting potential limitations of the current version of the database for these specific subclasses. Notably, while the deviations are insignificant for the overall classification as an ether lipid in general, they are still pronounced enough to prevent precise determination of the radyl binding type. By constructing simple linear models to describe the CCS and RT behavior of 1-acyl, 1-O-alkyl, and 1-O-alkenyl lipids, we demonstrate that these properties can be predicted based on basic lipid species information (Figure 6), paving the way for an implementation into generalized models. Notably, the linear model based on CCS values exhibited an overall higher error rate compared to the RT model. In connection with the above discussed additional influence of the precise molecular distribution of double bonds and carbon atoms on the CCS dimension, it follows that RT remains the more easily accessible parameter, while CCS captures more of the structural nuances. Furthermore, the model demonstrated a certain amount of orthogonality in the impact of different terms on CCS and RT. While the carbon chain length played a dominant role in the CCS model, the double bond term was most influential in the RT model (Figure 6).

### Future directions and technological advancements

IMS is an old concept^66^ that is currently evolving rapidly. This includes for example the maximal achievable resolution, the technical capabilities to apply the technology without significant ion losses^67^, but also the development of novel acquisition strategies^68^. As a result, the application of ion mobility is gaining increasing importance, particularly in complex lipid mixture analysis^69^, biomarker discovery^70,71^, structural biology^72^, and the isomer analysis within clinical science^73^. The CCS resolutions achieved in this study were not sufficient for baseline-separated differentiation of different isobaric ether lipid species (Figure 4). However, if current rapid technological developments continue, this point could be reached relatively soon. For the differentiation between 1-*O*-alkyl and 1-*O*-alkenyl lipids a three-fold improvement in IMS resolution would reduce the identification error rate below 5%, even without additional chromatographic separation (Figure 7). From that point, only a further two-fold increase would be needed to achieve baseline separation. It is already theoretically possible to achieve substantially higher resolutions by focusing on a very specific ion mobility range and allowing for long separation times. However, the necessary instrument parameters are then no longer compatible with more broadly applicable lipidomics methods. Despite current limitations, ion mobility already aids identification by providing clearer MS^2^ spectra and helping to rank the likelihood of 1-*O*-alkyl and 1-*O*-alkenyl forms and even modest improvements can significantly boost trust in this fourth dimension.

In summary, we demonstrate that IMS significantly enhances the differentiation between 1-*O*-alkyl and 1-*O*-alkenyl subspecies. Although current IMS capabilities alone are not yet fully sufficient to allow for univocal identification, combining IMS with orthogonal information such as RT greatly improves the reliability of lipid annotation.

## Experimental section

Methods are briefly summarized here, and a more detailed description can be found in Supplementary Text 2 and Supplementary Text 3:

Mouse tissues of wild type and *Peds1*-deficient mice were harvested at an age of 3-4 months, snap frozen in liquid nitrogen, and stored at −80°C. Animal breeding practices were approved by the Austrian Federal Ministry of Education, Science and Research (BMBWF-66.011/0100-V/3b/2019 and 2024-0.307.678).

Samples were subjected to lipid extraction according to Folch et al.^74^, stored at −20°C and resolved to obtain samples ready for LC-IM-MS measurements. Samples were measured in negative ESI mode on an Elute uHPLC System (Bruker Daltonics, Bremen, Germany) coupled with a TimsTOF Pro (Bruker Daltonics, Bremen, Germany), operated in imeX Ultra mode. Different Parallel Accumulation–Serial Fragmentation (PASEF) modes have been shown to be applicable for lipidomics^75^ and in the case of this study we have used data dependent acquisition-PASEF. Visual data inspection was performed using Data Analysis 5.3. (Bruker Daltonics, Bremen, Germany). A detailed description was included as Supplementary Text 2, for graphical visualization see Supplementary Figure 8.

Data extraction was done using an inhouse R pipeline (Supplementary Text 3) utilizing timsr^76^. For IMS calibration reference values were obtained manually from CCSbase (www.CCSbase.net/query_lipids), and a per sample linear model based on 1-acyl PE lipids fitted and applied to end up with comparable CCS values (Supplementary Figure 9). m/z values were not corrected. Retention times were aligned in a linear manner (Supplementary Figure 10). Database entries on molecular lipid species level matching our measured and annotated lipids by lm_id were extracted from lipidCCS, averaged to lipid species level, and compared with our results. Raw data, codebase and additional files are provided as Supplementary Dataset deposited at ZENODO (DOI: 10.5281/zenodo.11143478), and the lipidomics reporting checklist^77^ provided (DOI: 10.5281/zenodo.13963972).

## Author Contributions

J.K. and M.A.K. conceived and designed the study. J.K. developed the methodology. K.W. and M.A.K. provided resources. J.K. and K.L. performed experimental work. J.K. and L.N. analyzed data. M.L.F.-Q performed force field optimizations of gas phase structures. J.K. M.L.F.-Q and L.N. visualized results. J.K. wrote the original draft with input from K.W., L.N. and M.A.K.. All authors revised and agreed on the content of the manuscript.

## Acknowledgements

We thank Tereza Vlasáková for expert technical help. We also thank the Wellcome Trust Sanger Institute Mouse Genetics Project (Sanger MPG) and its funders for providing the mutant mouse line Tmem189tm1a(KOMP)Wtsi and INFRAFRONTIER/EMMA (https://www.infrafrontier.eu/). We acknowledge EuroHPC Joint Undertaking for awarding us access to MeluXina, Luxembourg. J.K. was funded by a DOC fellowship of the Austrian Academy of Sciences. This research was funded in part by the Austrian Science Fund (FWF) [10.55776/P33333, 10.55776/P34574] to M.A.K, FWF [10.55776/FG15] to L.N., K.W., and M.A.K., and FWF [10.55776/P34723] to K.W. M.L.F.-Q. is supported by the APART-MINT PostDoc fellowship of the Austrian Academy of Sciences (No. 11985).

## Notes

https://doi.org/10.5281/zenodo.13963971

### Competing Interest Statement

The authors have declared no competing interest.

