## Supplemental Material for "Trapped ion mobility spectrometry-guided molecular discrimination between plasmalogens and other ether lipids in lipidomics experiments"

Supplementary Figures

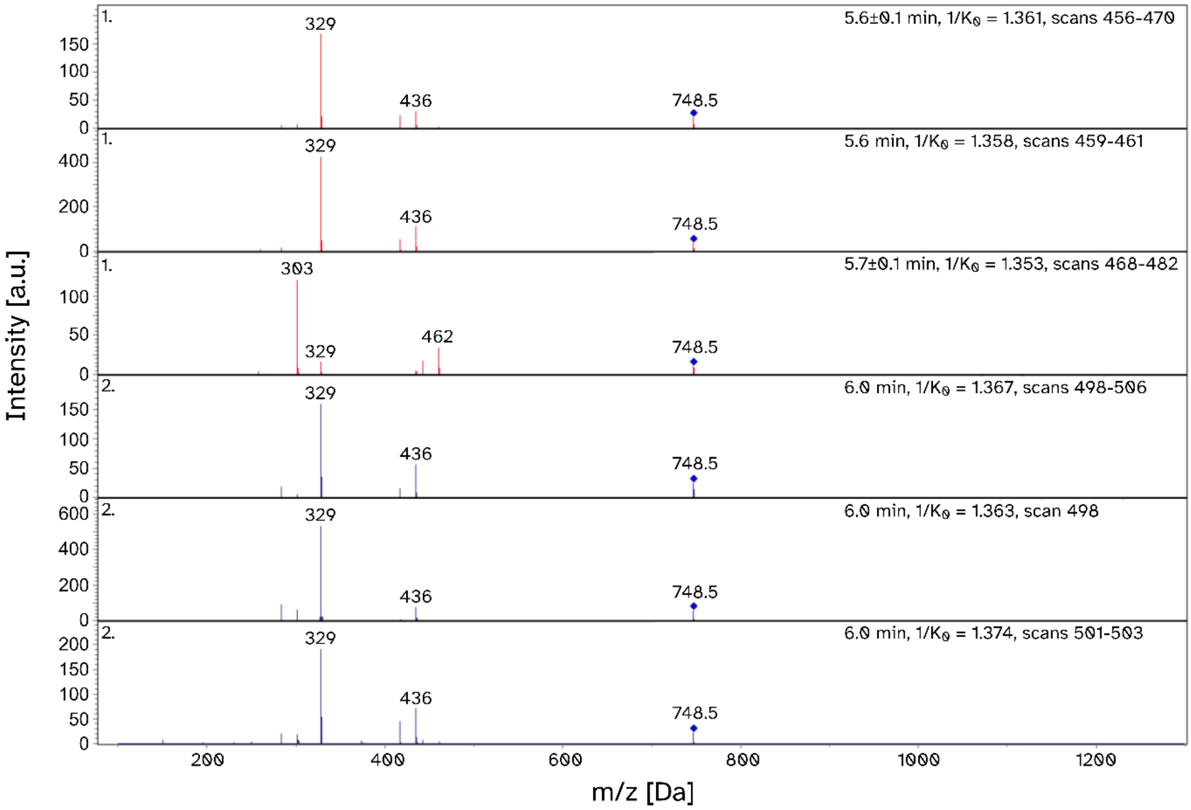

*Supplementary Figure 1: Example of MS$^2$ validation. m/z value plotted at the x-axis (Da) versus intensity (arbitrary units) as y-axis. The IMS dissected spectra for the first (red, ~5.6 min) and second (blue, ~ 6 min) region identified as PE P-38:5 (748.52 Da), from sample 301_H. MS$^2$ spectra of features annotated at molecular species level, which explain fatty acyl (FA) substitution at the sn-2 position, from which, by combination, the sn-1 fatty radyl residue was then deducted. m/z 329: FA 22:5; m/z 303: FA 20:4;*

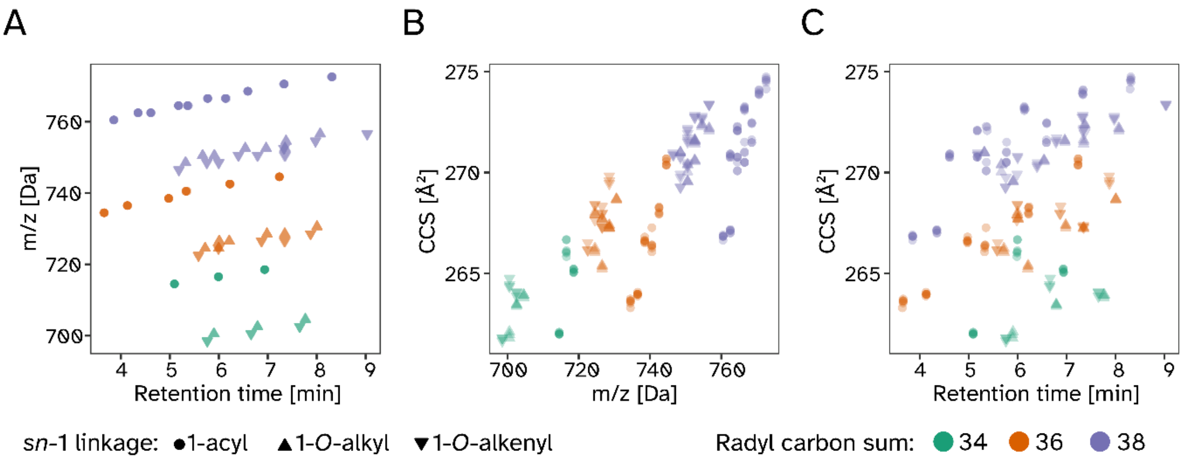

*Supplementary Figure 2: The extracted data of mouse* ***cerebrum*** *homogenates is displayed for all combined lipid features of wild type and* Peds1*-deficient samples (each n = 3). A) Retention time versus mass-over-charge plane. B) Mass-over-charge against ion mobility (displayed as collisional cross sections (CCS)) plane. C) The retention time versus ion mobility plane.*

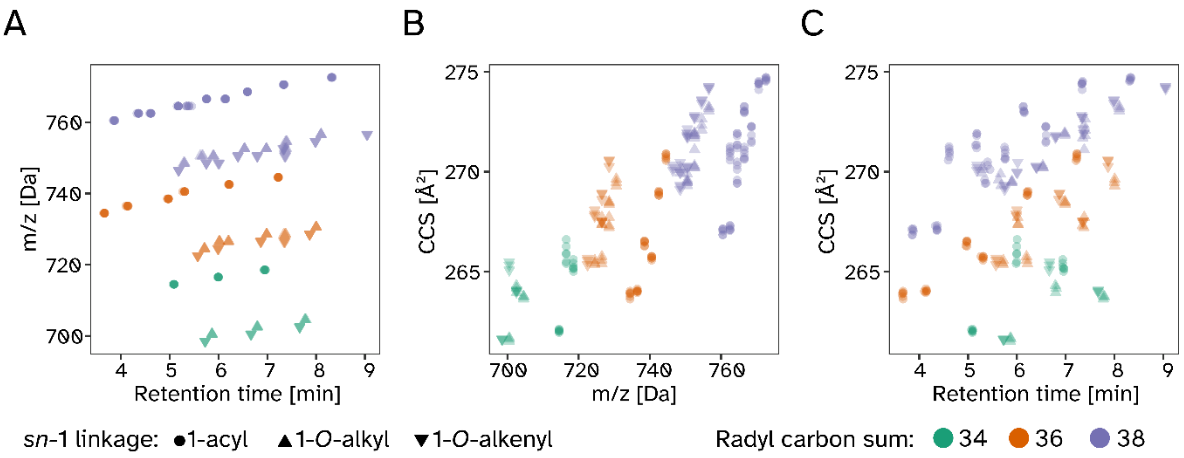

*Supplementary Figure 3: The extracted data of mouse* ***cerebellum*** *homogenates is displayed for all combined lipid features of wild type and* Peds1*-deficient samples (each n = 3). A) Retention time versus mass-over-charge plane. B) Mass-over-charge against ion mobility (displayed as collisional cross sections (CCS)) plane. C) Retention time versus ion mobility plane.*

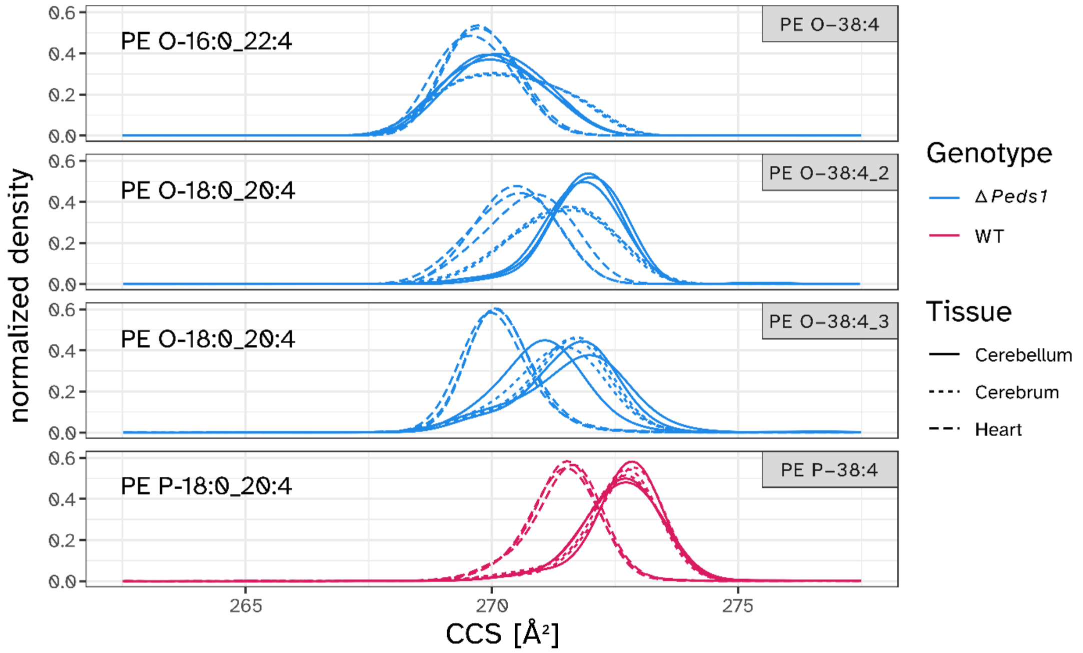

*Supplementary Figure 4: Individual CCS traces for selected isomeric species. CCS traces of fitted densities of individual features (long dashed line: heart, dashed: cerebrum, line: cerebellum). The four panels show different RT-m/z extraction windows, which were assigned to the stated molecular species level IDs on the left, while the lipid species level on the right give the standard feature identifiers (grey boxes) of this work.*

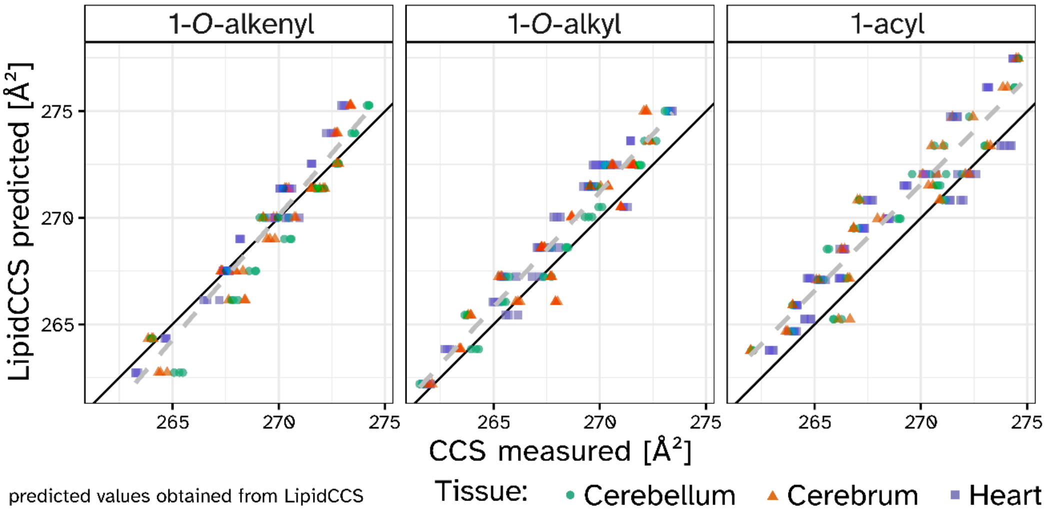

*Supplementary Figure 5: Database accuracy assessment. XY-scatter plots of predicted CCS values compared with the here measured values faceted by sn-1 linkage type into 1-acyl, 1-O-alkyl, and 1-O-alkenyl. The black line was drawn at a slope of 1, while the grey dashed line resulted from a linear model fit (no fixed intercept). The parallel offset in the 1-acyl panel indicates slight differences in the calibration regime. Symbols were drawn at a reduced alpha value and colored according to tissue of origin (circle: cerebellum, triangle: cerebrum, square: heart).*

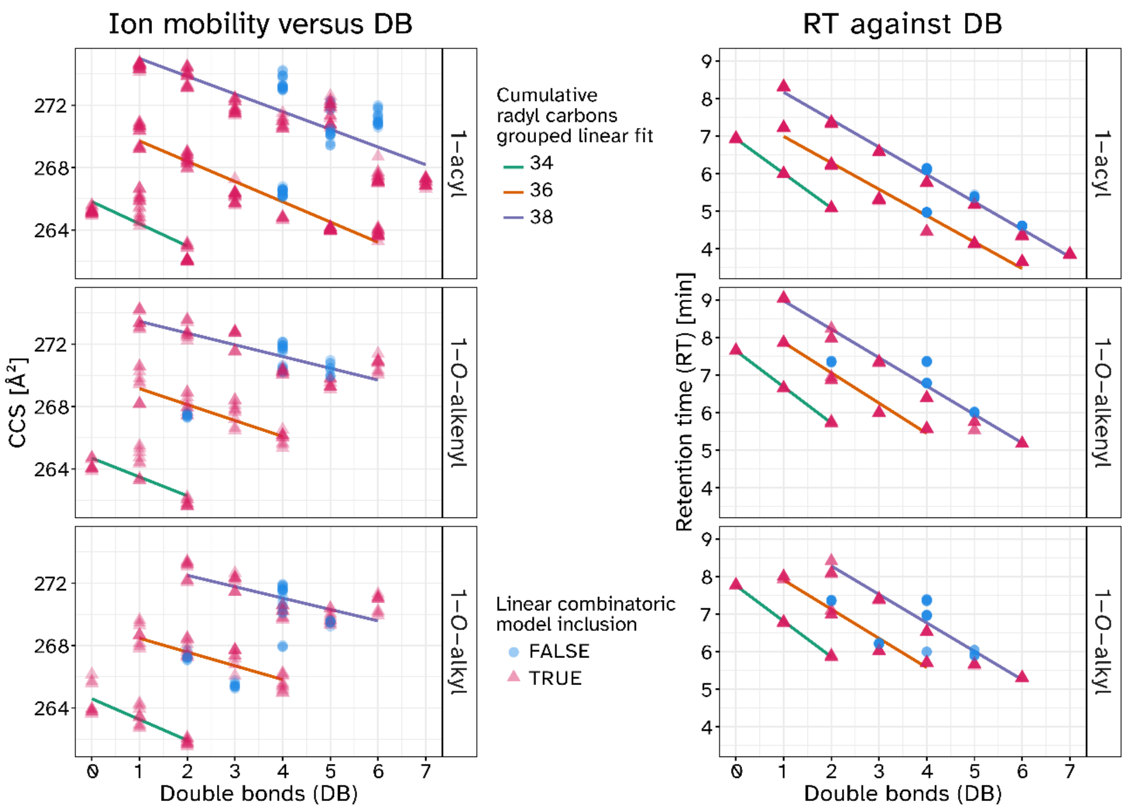

*Supplementary Figure 6: Plots of CCS and RT values against cumulative radyl double bonds stratified by linkage types. Linear regression lines per cumulative carbon atoms in the radyl side chains (CC) were plotted for 34 CC (red), 36 CCs (orange), and 38 CCs (green). The color of the data points indicates whether the statistical analysis of the regression model would indicate them as within the deviation of the model (red) or as outliers (blue). Observe that while we used a model with a larger set of free parameters, i.e. different inclinations for different CC, with respect to Figure 6 this leads to only minor improvement of the data fit.*

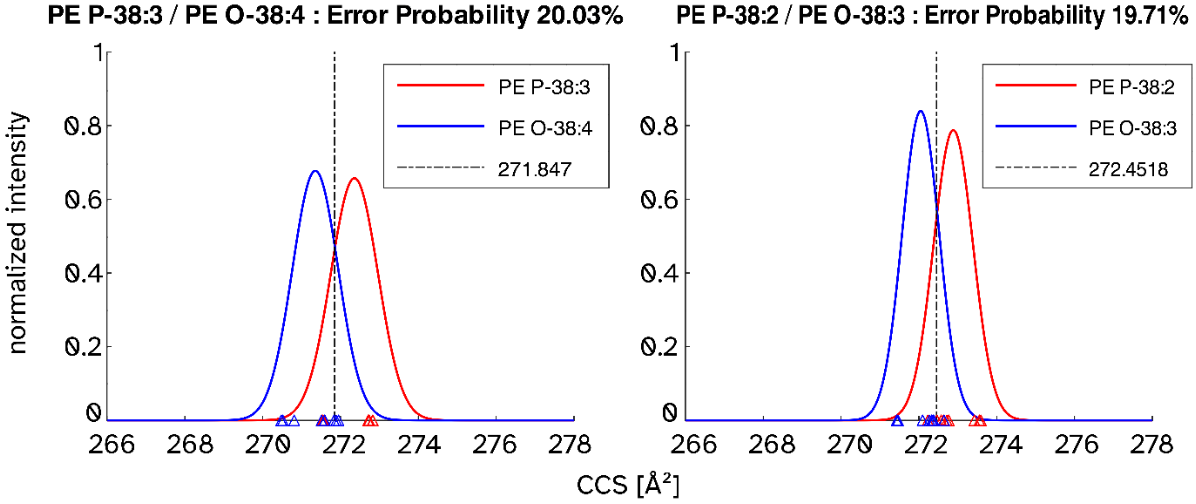

*Supplementary Figure 7: CCS versus normalized intensity plots for two selected lipid pairs: 1-O-alkenyl (red) and 1-O-alkyl (blue). A dashed vertical line marked the midpoint of the overlap, at which the given error probability applied. Plotted on the left for the PE P-38:3 / PE O-38:4 pair, and on the right side for the PE P-38:2 / PE O-38:3 pair. Individual measurements are depicted as colored triangles.*

### Supplementary Text 1 - Theoretical prediction of mobility values

We predicted individual CCS values for abundant 1-acyl, 1-*O*-alkyl, and 1-*O*-alkenyl phosphatidylethanolamine species. From the multitude of different combinations of radyl chains, we selected three pairs with palmitoleyl or oleyl residues linked at the *sn*-1 position, and either docosahexaenoyl or arachidonoyl linked at the *sn*-2 position. This was done in a manner that ensured that the cumulative side chain characteristics were kept constant (18:0/20:4 and 16:0/22:4), and only the sn-1 linkage type varied. For each lipid species .mol files were prepared and subjected to a energy minimization in the gas phase using the AMBER10:EHT forcefield implemented in MOE (Molecular Operating Environment (MOE), 2022.02 Chemical Computing Group ULC, 910-1010 Sherbrooke St. W., Montreal, QC H3A 2R7). The obtained structures were reported as .pdb files and are included in the Supplementary Dataset (DOI:10.5281/zenodo.11143478).

To obtain CCS values from the minimized structures, the pdb files, in $[M\text{-}H]^-$ form, were converted into .xyz files using openbabel GUI (3.1.1)[1], and processed via the Projected Superposition Approximation (PSA) webserver provided by the Bleiholder laboratory at FSU (http://psa.chem.fsu.edu/)[2], with the following input parameters kept constant for the six different files processed: task psa.task, psa_general_buffergas nitrogen, psa_general_temperature 300, psa_projection_accuracy 0.01, psa_projection_integration_accuracy 0.009, psa_shape_accuracy 0.01, psa_shape_maxiter 25, psa_shape_meshfactor 1; For one file (new_LMGP02010116.xyz) the psa_general_temperature was varied from 300 – 500 by 20 K steps, to estimate the temperature parameter influence. We found that with increasing temperature the resulting CCS values decreased (300 K: 418.0 Å² to 500 K: 359.7 Å²). Comparing the PSA predicted values with their respective measured counterparts we observed clear discrepancies. Measured ether lipid CCS traces were visualized in Supplementary Figure 4.

The effect sizes observed in the PSA results were substantially higher, compared to measured values (Supplementary Table 1 *versus* Supplementary Table 2). When analysed for their qualitative behavior and disregarding numeric differences, we found that the PSA values of two out of three lipid species pairs in respect to their measured values showed the same trend: in a subclass specific view 1-acyl presented with the lowest values, while for 1-*O*-alkyl and 1-*O*-alkenyl values increased. 1-*O*-acyl and 1-*O*-alkenyl pairs containing a FA 22:4 side chain had lower CCS values compared with their FA 20:4 containing isobars in both, predicted as well as measured results (Supplementary Table 2).

### Supplementary Text 2 - From living animals to measured data

Tissue homogenization, and lipid extraction was performed as previously described[3].

#### *Generation and tissue collection of Peds1-deficient mice*

This study used mice with a deficiency in the plasmanylethanolamine desaturase (PEDS1) enzyme (Tmem189tm1a(KOMP)Wtsi, obtained from the Wellcome Sanger Institute, Hinxton, Cambridge, UK,[4]) and wild type littermate controls (all on C57bl/6N background). Animal breeding was approved by the Austrian Federal Ministry of Education, Science and Research (BMBWF-66.011/0100-V/3b/2019 and 2024-0.307.678). Breeding, housing and genotyping followed established protocols[5]. Briefly, mice lived in individually ventilated cages with nesting material under a 12-hour light/dark cycle. They received standard chow (Sniff Spezialdiäten GmbH, standard chow V1534-300, inhouse autoclaved) and water freely. Tissues were collected from 3-4 months old female and male *Peds1*-deficient mice and their wild type littermates. Animals were sacrificed by cervical dislocation, harvested tissues were snap-frozen in liquid nitrogen and stored at -80°C for further analysis.

#### *Tissue homogenization and analyte isolation*

We homogenized the harvested mouse tissues in 1x PBS with an Ultra-Turrax (T10, IKA, Staufen, Germany). After additional shearing in the presence of glass beads using a MM400 mixer mill (IKA, Germany) the protein content was determined with a standard Bradford assay, calibrated with a BSA dilution series. 300 µg total protein aliquots were extracted according to the Folch method[6]. Lipid extracts were dried under a $N_2$-flow and stored at -20°C until reconstitution in 100 µL of 62% B running eluent mixture (see below) prior to measurement.

#### *LC-IM-MS/MS analysis*

Lipidomic analysis was performed on a Bruker Elute HPLC system coupled to a timsTOF Pro (Bruker Daltonics, Bremen, Germany). 10 µL of each sample was injected at a flowrate of 0.4 mL/min onto a temperature-controlled (50 °C) Agilent Poroshell 120 EC-C8 2.7mm 2.1x100mm column (Agilent Technologies, Santa Clara, USA). Chromatographic separation was achieved with a reversed phase gradient using the running eluents A (4/6 $H_2O$/acetonitrile (v:v), 10 mM ammonium formate, 0.2% formic acid) and B (9/1 isopropanol/acetonitrile (v:v), 10 mM ammonium formate, 0.2% formic acid). These were prepared freshly before measurement and sonicated for at least 30 minutes until no ammonium formate remains were visible. HPLC starting conditions were 56% A, which was maintained in isocratic flow for 2 minutes, followed by a linear decrease to 35% A over 11 minutes. Next, buffer A content was reduced to 3% while increasing the flow rate to 0.6 µL/min over two minutes. This was followed by a gradient to 1% A until minute 19 at the high flow rate. Within the next 1-minute period, flowrate and A content were reset to starting conditions, and the column was equilibrated further for another 3.5 minutes. The maximum pressure was set to 650 bar, and the total method runtime was 23.5 minutes. The autosampler was operated in µL pickup mode, with sample speed medium, no air segments, a sample environment temperature of 10°C, and a needle height of 2 mm. The wash-phases were programmed to wash with 1000 µL 90% IPA (Solvent 2), 1000 µL 50% ACN (Solvent 1), 1000 µL 90% IPA (Solvent 2), and 1500 µL 50% ACN (Solvent 1).

Analytes were ionized via the Apollo II source (ESI), with a dry temperature of 250°C, end plate offset 500 V, nebulizer N2-pressure 3.3 bar, capillary voltage 4500 V and dry gas flow of 10 L/min. The mass spectrometer was operated in negative ion polarity within the mass range of 100 - 1640 m/z. The TIMS cell was set to cover an inverse ion mobility ($1/K_0$) range of 1.20 - 1.45 Vs/cm² in imeX Ultra mode at maximum resolution, which resulted in a ramp time of 714.9 ms and a spectra rate of 1.39 Hz. In comparison to other methods, these were very narrow IMS ramp settings, which were tailored to phosphatidylethanolamine lipids. The accumulation setting was locked to the selected mobility range, and the 100% duty cycle setting activated. The TIMS settings were changed to cover the IMS calibrants range during injection (17.5 – 17.7 min) of 20 µL of Agilent Tuning mix low conc. (Agilent Technologies, Santa Clara, USA) diluted 1:20 delivered from a syringe pump (10 mL Hamilton syringe, otofControl settings adjusted accordingly).

***Instrument calibration and tuning***

The mass detector was calibrated linearly using a manual selection of ions from the Agilent Tuning mix low conc. [m/z: 431.9823; 601.9788; 1033.9885; 1433.9617; 1633.9501]. Prior to the actual TIMS calibration the $N_2$-Flowrate was adjusted accordingly to the instrument guidelines (IIa. Getting Started: ESI-source with Ion Mobility – Revision B (October 2019) and set to 132+-1eV for the m/z 622.0289 calibration mix component. The TIMS dimension was calibrated linearly prior to measurement using three selected ions from the Agilent Tuning mix low conc. [*m/z*, $1/K_0$: (601.9790, 0.8784 Vs cm$^{-2}$), (1033.9881, 1.2502 Vs cm$^{-2}$), (1333.9689, 1.4038 Vs cm$^{-2}$)] in manual mode (imeX Ultra).

The Collision cell parameters were set to Collision Energy: 10.0 eV, Collision RF: 1200.0 Vpp, Transfer Time: 80.0 µs, and Pre Pulse Storage: 15.0 µs. The quadrupole was set to Ion Energy: 5.0 eV and Low Mass: 150.00 m/z. Transfer parameters were set to Deflection 1 Delta: -80.0 V, Funnel 1 RF: 250.0 Vpp, Funnel 2 RF: 250.0 Vpp, isCID Energy: 0.0 eV, and Multipole RF: 200.0 Vpp. For further method details see the method and tuning report included in the Supplementary Dataset (DOI:10.5281/zenodo.11143478).

### Supplementary Text 3 – Data extraction, modeling and visualization

Data analysis was done primarily in R (see Supplementary Dataset, where all the herein referenced files are provided, DOI:10.5281/zenodo.11143478), and organized in such, that all analyses were described in the respective quarto (.qmd) files located within a qmd folder, functions and other larger code snippets are saved in corresponding .R files located in the R folder (automatically sourced in their parent files). Data was saved into the data folder, where data/data_raw contains the measured bruker .d files, while data_processed contains important tables written to disk in .Rdata format. Furthermore, the CCS data from lipidCCS can be found in CCS_data while _CCS_data contains the files uploaded to predict CCS values for.

***Data integration***

The inhouse data analysis pipeline sketched in Supplementary Figure 8 was setup in R, running in a linux based docker container. R (4.3.0), Rstudio (2023.03.1 Build 446) and the following packages were utilized throughout the course of data extraction from the raw data files: broom (1.0.4), devtools (2.4.5), ggExtra (0.10.0), here (1.0.1), tidyverse(2.0.0), timsr (0.0.3), latex2exp (0.9.6), patchwork (1.1.2).

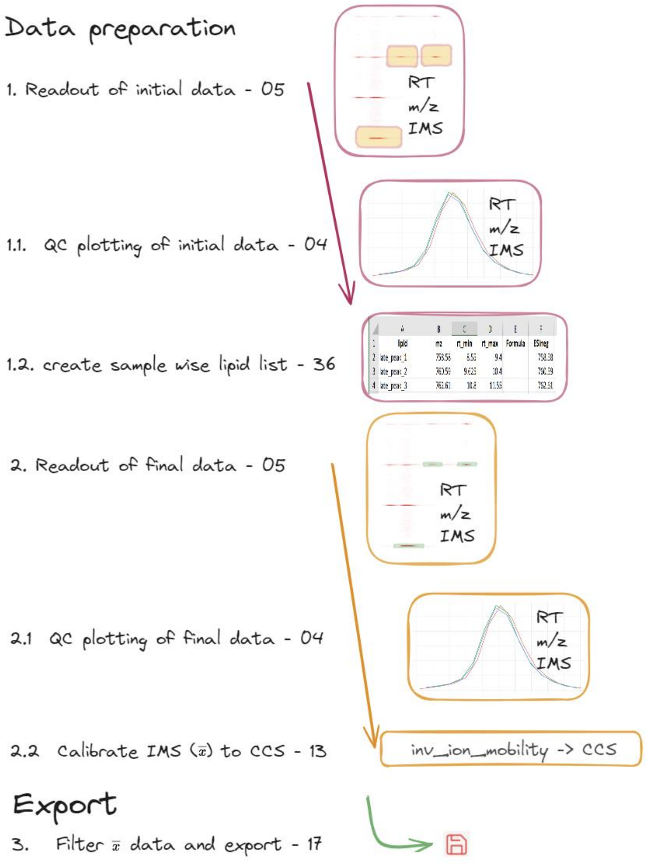

*Supplementary Figure 8: Schematic representation of data extraction steps.*

Group wise integration lists were generated and manually optimized (containing lipid, m/z as measured, retention time (RT) start & end, Formula, and calculated m/z information). The optimized integration list is provided within the Supplementary Dataset (DOI:10.5281/zenodo.11143478, Peaklist_8.xlsx). With a mass tolerance of 60 mDa, data extraction of the monoisotopic peak was performed (05_Readout_PLOP4_manual.qmd). The resulting data set was then visualized for graphical quality control (04_JK_Control-plotting_RT-and-IMS.qmd), and from the dataset the optimal m/z values for data extraction were determined and a sample-wise integration list was created (Peaklist_9.xlsx). This list was used for the conclusive data extraction (05_Readout_PLOP4_manual.qmd) with a narrow tolerance of 15 mDa and the data plotted for additional graphical verification of data integrity (04_JK_Control-plotting_RT-and-IMS.qmd). The weighted.mean masses of the extracted PE lipids were found to deviate from the theoretical ones by around -7 mDa. This dataset was summarized to obtain mean values for RT, m/z, IMS, and an intensity sum (area).

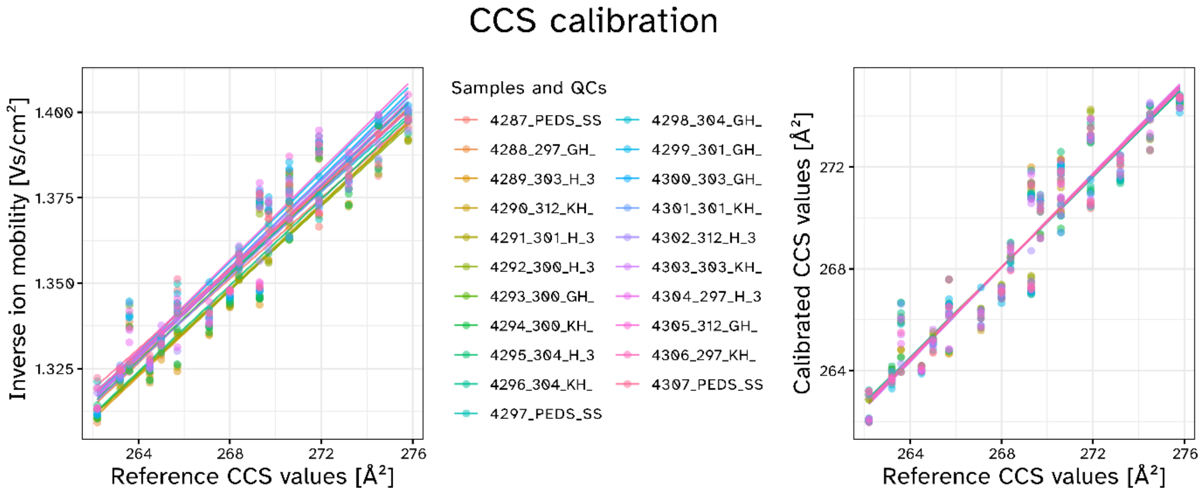

*Supplementary Figure 9: CCS calibration. Left: Averaged IMS values linear models per sample for 1-acyl lipids, plotting inverse ion mobility versus predicted database reference CCS values (x-axis). Right: measured and calibrated CCS values (y-axis, Å²) plotted against the used reference CCS values (CCS_ref, x-axis, Å²). Sample nomenclature in the figure legend according to <injection number>_<mouse id>_<tissue id>_…; with tissue id: cerebrum (GH), cerebellum (KH), heart (H);*

Next, a sample wise ion mobility calibration of the collision cross section (CCS) values was performed with reference values obtained from CCSbase (www.ccsbase.net/lipids_query, last accessed 2024-05-02; 2023-02-24_JK_CCSbase-manual-extraction-on-our-level.xlsx, Supplementary Figure 9) followed by a linear RT correction (13_JK_CCS_corrections.qmd). For the CCS and RT dimension no trend in relation to sample order during measurement was observed. Only in one instance a linear RT correction seemed necessary (as visualized in Supplementary Figure 10), to correct for a pressure glitch during data acquisition, but of course an identical correction was applied to all individual samples.

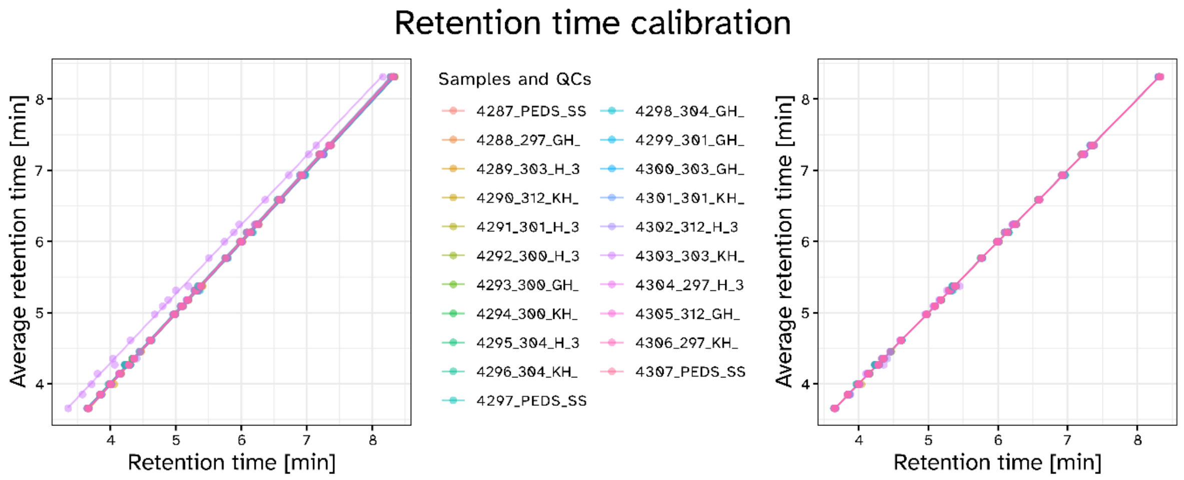

*Supplementary Figure 10: Linear retention time (RT) correction of measured values. Left panel shows data before RT calibration, right panel after RT calibration. This clearly evens out the observed RT shift due to a pressure glitch in Sample 303_KH. Mean RT [min] across all samples and analytes plotted against the individual RT [min] on the x-axis. Sample nomenclature in the figure legend according to <injection number>_<mouse id>_<tissue id>_…; with tissue id: cerebrum (GH), cerebellum (KH), heart (H);*

Ambiguous features were removed, and the resulting data exported in a full (2023-09-25_data_all_PLOP2_LNeumann.csv == **data_all**) and filtered for the highest distinct lipid species level observations (2023-09-25_data_PLOP2.csv == **data**) version (17_JK_Datenexport.qmd, also containing a global data dictionary). A tissue averaged table of all relevant PE peaks analyzed in this study was provided as overview_table.xlsx within the Supplementary Dataset (DOI:10.5281/zenodo.11143478, see: _output\19_JK_overview_dataframe-creation\ overview_table.xlsx).

***Data modeling and visualization***

A windows based R (4.3.2) installation operated through Rstudio (2023.12.1 Build 402), and the following packages was utilized for graphical visualization of the integrated and calibrated data set: tidyverse (2.0.0), broom (1.0.5),rgoslin (1.3.1), enviPat (2.6),extrafont (0.19), extrafontdb (1.0), ggblend (0.1.0), ggforce (0.4.1), ggrepel (0.9.3), ggtext (0.1.2), lipidmapsR (1.0.4), patchwork (1.2.0), RcppRoll (0.3.0).

For Figure 2 RAW data visualizations were exported from the Data analysis software (Version 5.3. build 236.352.5870 64-bit, Bruker Daltonics, Bremen, Germany) with the following parameters: RT range 2-10 min, m/z range 745-755 Da, and IMS range 1.34-1.39 Vs/cm², mean values (39P_JK_dots-for-figure-2-positioning.qmd) were superimposed. Figure 3 was created solely in R using **data_all** (38P_JK_Figure3ABC-pannels.qmd). Part A of Figure 4 was created by 41P_JK_PE-O-38-4_P-38-3-traces-Figure4.qmd and the statistical assessment was done as described in 02_JK_RT-and-IMS-PLOP1-mean-plots.qmd. Figure 5 was created in script 20P_JK_measured_vs_predicted_data.qmd using **data_all**. Figure 6 was created from fitted results in R (A: 38P_JK_Figure3ABC-pannels.qmd, B: 32P_JK_CCS-model-plotting.qmd).

For linear model fitting we used **data**, and fitted a linear model, in respect to CC, DB, and *sn*-1 linkage type for RT and CCS independently. The error contributions of each model were normalized to their total sum and then visualized. The actual fit and ANOVA table outputs can also be found in 32P_JK_CCS-model-plotting.qmd. The code used to create Figure 7 from data_all can be found in 40P_JK_Resolution-vs-Error-plotting.qmd. All main figures were finalized in Affinity Designer 1.10 (SansSerif, Nottingham, United Kingdom). All R code files and additional files were deposited at Zenodo (10.5281/zenodo.11143478).

### Supplementary Tables

*Supplementary Table 1: PSA calculation for a set of selected lipid in their protonated $[M-H]^-$ form.*

| **Radyl-substi-tution** | ***lm_id*** | ***T[K]*** | ***PA [Å²]*** | ***RHO*** | ***PSA [Å²]*** | ***error [Å²]*** |
|---|---|---|---|---|---|---|
| *16:0_22:4* | LMGP02010116 | 300 | 409 | 1.022 | 418 | 3.8 |
| *18:0_20:4* | LMGP02010118 | 300 | 414 | 1.02 | 422.2 | 3.2 |
| *O-16:0_22:4* | LMGP02020037 | 300 | 397.2 | 1.02 | 405.1 | 3.5 |
| *O-18:0_20:4* | LMGP02020092 | 300 | 370.9 | 1.032 | 382.9 | 3 |
| *P-16:0_22:4* | LMGP02030033 | 300 | 344.2 | 1.044 | 359.2 | 2.9 |
| *P-18:0_20:4* | LMGP02030003 | 300 | 380.5 | 1.029 | 391.6 | 3.9 |

*Supplementary Table 2: Measured CCS values of the same lipids as listed in Supplementary Table 1. The rows formatted in bold indicate the most intense entries found per substitution group.*

| lipid | Inverse ion mobility. [Vs cm⁻²] | Intensity [a.u.] | Area [a.u.] | RT [min] | CCS [Å²] |
|---|---|---|---|---|---|
| PE_38:4 | 1.37266161 | 173.5 | 49916 | 5.76 | 270.8 |
| **PE_38:4_2** | **1.38991441** | **2756.9** | **3025412** | **6.13** | **273.4** |
| PE_O-38:4 | 1.36739648 | 387.9 | 414403 | 6.53 | 270.1 |
| **PE_O-38:4_2** | **1.37574172** | **752.8** | **519394** | **6.96** | **271.4** |
| PE_O-38:4_3 | 1.37176286 | 151.1 | 36669 | 7.38 | 270.8 |
| PE_P-38:4 | 1.36749135 | 310.2 | 222341 | 6.40 | 270.2 |
| **PE_P-38:4_2** | **1.37491329** | **731.3** | **515569** | **6.79** | **271.3** |
| PE_P-38:4_3 | 1.37611802 | 103.0 | 16409 | 7.36 | 271.5 |

(4) White, J. K.; Gerdin, A.-K.; Karp, N. A.; Ryder, E.; Buljan, M.; Bussell, J. N.; Salisbury, J.; Clare, S.; Ingham, N. J.; Podrini, C.; Houghton, R.; Estabel, J.; Bottomley, J. R.; Melvin, D. G.; Sunter, D.; Adams, N. C.; Tannahill, D.; Logan, D. W.; MacArthur, D. G.; Flint, J.; Mahajan, V. B.; Tsang, S. H.; Smyth, I.; Watt, F. M.; Skarnes, W. C.; Dougan, G.; Adams, D. J.; Ramirez-Solis, R.; Bradley, A.; Steel, K. P.; Baker, L.; Barnes, C.; Beveridge, R.; Cambridge, E.; Carragher, D.; Chana, P.; Clarke, K.; Hooks, Y.; Igosheva, N.; Ismail, O.; Jackson, H.; Kane, L.; Lacey, R.; Lafont, D. T.; Lucas, M.; Maguire, S.; McGill, K.; McIntyre, R. E.; Messager, S.; Mottram, L.; Mulderrig, L.; Pearson, S.; Protheroe, H. J.; Roberson, L.-A.; Salsbury, G.; Sanderson, M.; Sanger, D.; Shannon, C.; Thompson, P. C.; Tuck, E.; Vancollie, V. E.; Brackenbury, L.; Bushell, W.; Cook, R.; Dalvi, P.; Gleeson, D.; Habib, B.; Hardy, M.; Liakath-Ali, K.; Miklejewska, E.; Price, S.; Sethi, D.; Trenchard, E.; von Schiller, D.; Vyas, S.; West, A. P.; Woodward, J.; Wynn, E.; Evans, A.; Gannon, D.; Griffiths, M.; Holroyd, S.; Iyer, V.; Kipp, C.; Lewis, M.; Li, W.; Oakley, D.; Richardson, D.; Smedley, D.; Agu, C.; Bryant, J.; Delaney, L.; Gueorguieva, N. I.; Tharagonnet, H.; Townsend, A. J.; Biggs, D.; Brown, E.; Collinson, A.; Dumeau, C.-E.; Grau, E.; Harrison, S.; Harrison, J.; Ingle, C. E.; Kundi, H.; Madich, A.; Mayhew, D.; Metcalf, T.; Newman, S.; Pass, J.; Pearson, L.; Reynolds, H.; Sinclair, C.; Wardle-Jones, H.; Woods, M.; Alexander, L.; Brown, T.; Flack, F.; Frost, C.; Griggs, N.; Hrnciarova, S.; Kirton, A.; McDermott, J.; Rogerson, C.; White, G.; Zielezinski, P.; DiTommaso, T.; Edwards, A.; Heath, E.; Mahajan, M. A.; Yalcin, B. Genome-Wide Generation and Systematic Phenotyping of Knockout Mice Reveals New Roles for Many Genes. *Cell* **2013**, *154* (2), 452–464. https://doi.org/10.1016/j.cell.2013.06.022.

(5) Werner, E. R.; Keller, M. A.; Sailer, S.; Lackner, K.; Koch, J.; Hermann, M.; Coassin, S.; Golderer, G.; Werner-Felmayer, G.; Zoeller, R. A.; Hulo, N.; Berger, J.; Watschinger, K. The TMEM189 Gene Encodes Plasmanylethanolamine Desaturase Which Introduces the Characteristic Vinyl Ether Double Bond into Plasmalogens. *Proc. Natl. Acad. Sci. U. S. A.* **2020**, *117* (14), 7792–7798. https://doi.org/10.1073/pnas.1917461117.

(6) Folch, J.; Lees, M.; Sloane Stanley, G. H. A Simple Method for the Isolation and Purification of Total Lipides from Animal Tissues. *J. Biol. Chem.* **1957**, *226* (1), 497–509.
